# Dominant negative effects of *SCN5A* missense variants

**DOI:** 10.1101/2021.09.22.461398

**Authors:** Matthew J. O’Neill, Ayesha Muhammad, Bian Li, Yuko Wada, Lynn Hall, Joseph F. Solus, Laura Short, Dan M. Roden, Andrew M. Glazer

## Abstract

**Introduction:** Up to 30% of patients with Brugada Syndrome (BrS) carry loss-of-function (LoF) variants in the cardiac sodium channel gene *SCN5A*. Recent studies have suggested that the *SCN5A* protein product Na_V_1.5 can form dimers and exert dominant negative effects.

**Methods:** We identified 35 LoF variants (<10% peak current compared to wild type (WT)) and 15 partial LoF variants (10-50% peak current compared to WT) that we assessed for dominant negative behavior. *SCN5A* variants were studied in HEK293T cells alone or in heterozygous co-expression with WT *SCN5A* using automated patch clamp. To assess clinical risk, we compared the prevalence of dominant negative vs. putative haploinsufficient (frameshift/splice site) variants in a BrS case consortium and the gnomAD population database.

**Results:** In heterozygous expression with WT, 32/35 LoF variants and 6/15 partial LoF showed reduction to <75% of WT-alone peak I_Na_, demonstrating a dominant negative effect. Carriers of dominant negative LoF missense variants had an enriched disease burden compared to putative haploinsufficient variant carriers (2.7-fold enrichment in BrS cases, p=0.019).

**Conclusions:** Most *SCN5A* missense LoF variants exert a dominant negative effect. Cohort analyses reveal that this class of variant confers an especially high burden of BrS.

## Introduction

Brugada Syndrome (BrS) is a clinical arrhythmia syndrome with characteristic EKG changes in the absence of underlying structural heart abnormalities (1). While often asymptomatic or clinically unrecognized, sudden cardiac death (SCD) due to ventricular tachyarrhythmia can be the sentinel manifestation. Up to 30% of BrS patients have heterozygous loss-of-function (LoF) variants in the cardiac sodium channel gene *SCN5A*, which encodes the channel protein Na_V_1.5 (2). A recent evaluation by ClinGen asserted that *SCN5A* was the only gene with strong evidence for Mendelian associations with BrS (3). LoF *SCN5A* variants are also associated with other arrhythmias including sick sinus syndrome (4) and progressive cardiac conduction disease (5).

Over 100 LoF variants within *SCN5A* have been reported across multiple variant classes including missense, nonsense, splice-altering, and frameshift/premature truncation (2, 6). *SCN5A* encodes a channel with 4 transmembrane domains, each consisting of 6 transmembrane segments (7). Na_V_1.5 has traditionally been thought to function as a monomer; however, a recent study indicated that Na_V_1.5 can form dimers with coupled intracellular trafficking and/or gating at the plasma membrane (8). Similar to variants in established multimeric proteins that can generate dominant negative effects, several missense *SCN5A* variants with dominant negative effects on trafficking or coupled gating at the cell surface have been reported *in vitro* and *in vivo* (9–11). However, the dominant negative behavior of most of the approximately 40 known LoF missense variants in *SCN5A* has not been tested. Moreover, the degree of dominant negative effects among partial LoF missense variants has not been evaluated.

Variable penetrance is a hallmark of pathogenic BrS variants, and the extent to which distinct pathogenic mechanisms (e.g., dominant negative vs haploinsufficiency) contribute to this effect is unknown. Large cohort studies and variant curation efforts provide datasets of *SCN5A* variants associated with BrS cases (2, 6, 12). In addition, large population cohorts such as gnomAD provide sets of individuals more likely representing putative controls (13). Together, these datasets enable the comparison of BrS disease risk among different variant classes.

Here, we study the prevalence of the dominant negative effect among *SCN5A* LoF and partial LoF missense variants. We use case and control cohorts to test the relative BrS disease risk of dominant negative missense variants compared with other variant classes.

## Methods

### Selection of Variants

Variants for this study were selected from previously published functionally characterized variants (6, 14). Variants with peak currents <10% compared to WT were considered LoF, and variants with peak currents between 10-50% compared to WT were considered partial LoF. A full list of variants in this study is presented in Supplementary Table 1.

### SCN5A Mutagenesis

The *SCN5A* variant plasmids were mutagenized using a previously described “zone” system (14). Briefly, *SCN5A* individual zones on small plasmids were mutagenized using the QuikChange Lightning Multi kit (Agilent) with primers designed using the online QuikChange Primer Design tool. Primers used in this study are listed in Supplementary Table 2. The variant-containing zone was then subcloned by restriction digestion into a plasmid containing an AttB:SCN5A:IRES:mCherry-blasticidinR plasmid (14–16). The entire sequence of the zone containing the variant was confirmed by Sanger sequencing. In a previous study of 82 variants generated by this approach, 0/82 plasmids had any additional *SCN5A* mutations outside the target zone (14). All analyses used the most common *SCN5A* transcript in the adult heart, including the adult isoform of exon 6 and a deletion of the alternatively spliced Gln1077 residue (ENST00000443581). As per convention, all variants are named in accordance with the full 2,016 amino acid form (ENST00000333535).

### Description of Cell Lines

All experiments used Human Embryonic Kidney HEK293T “negative selection” landing pad (LP) cells as previously described (gift of Kenneth Matreyek) (14–16). The AttB/AttP LP allows a single integration event per cell and a consistent level of target gene expression (Figure 1). Homozygous experiments were carried out in LP cells (Figure 1A). Plasmids carrying *SCN5A* variants were transfected along with transposase and integrated into the LP site to allow stable expression. We termed these lines LP-SCN5A.

**Figure 1.**
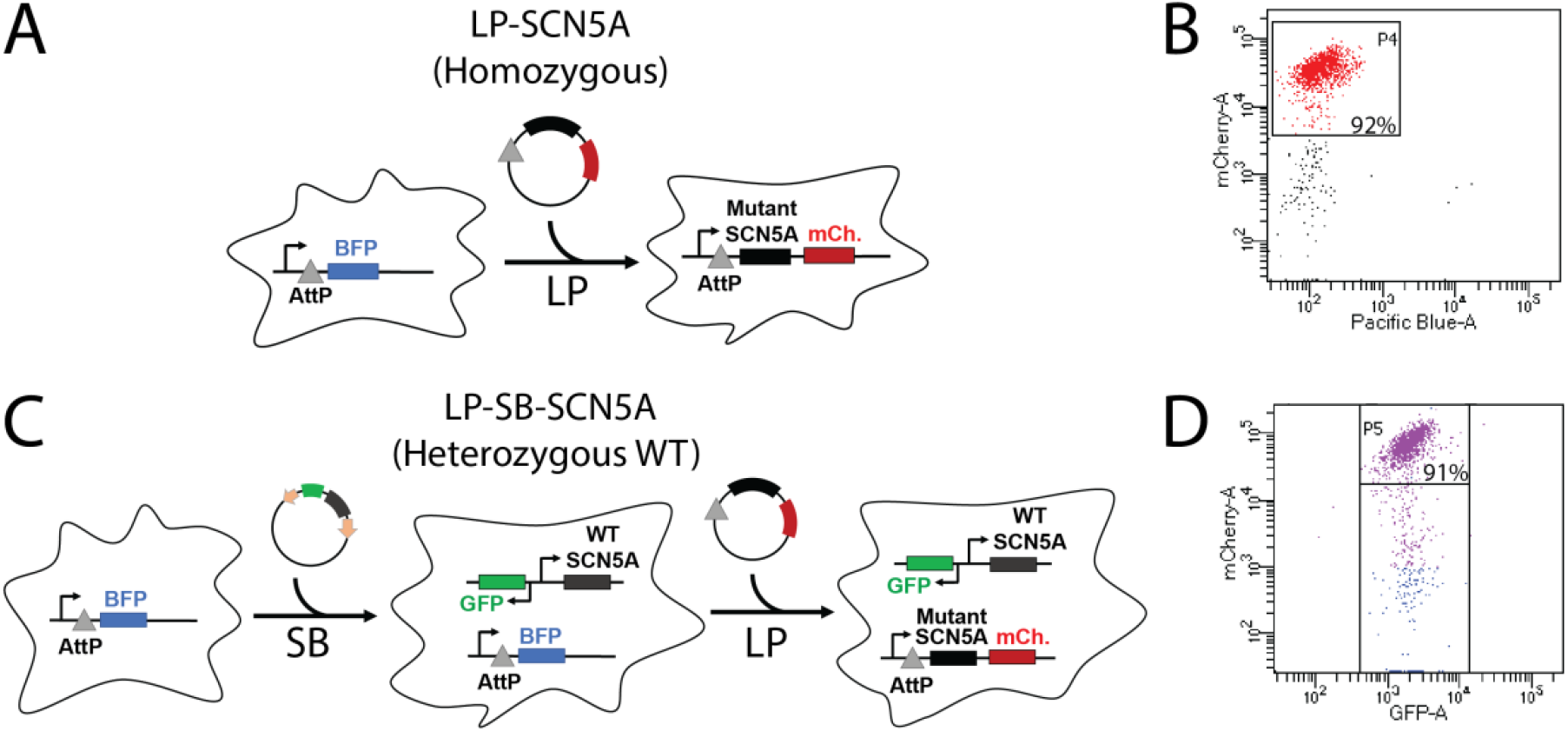
Stable cell lines used in this study. 1 or 2 copies of *SCN5A* were inserted into engineered HEK293 LP cells. The Landing Pad (LP) comprises an AttP and BFP locus, and allows insertion of a single insert per cell. A second Sleeping Beauty (SB) transposon system was used to introduce a second copy of the gene for heterozygous experiments. **A**. Design of homozygous LP-SCN5A cell line with LP integration. **B.** Analytical flow cytometry after incorporation of plasmid into the LP. Cells that do not have BFP expression and highly express mCherry (P4 gate) have a successful integration. **C**. For heterozygous experiments, we used a combination of LP and SB systems. First, a SB plasmid bearing a WT copy of *SCN5A* was randomly inserted into the genome. A clone of these cells was identified that has an equal level of Na_V_1.5 in patch clamp experiments to typical LP expression (Figure 2). Next, a second copy of *SCN5A* bearing WT or variant was incorporated through the LP system. **D.** Results of flow cytometry after SP and LP integration. Cells express GFP associated with SB integration, and mCherry after LP integration (P5 gate).

For heterozygous expression, we first generated LP cells stably expressing WT *SCN5A* in a non-LP site using the Sleeping Beauty (SB) transposon system and identified a clone of this cell line with peak sodium current (I_Na_) equivalent to that observed with WT *SCN5A* in the LP site (Figure 1B). We then generated cell lines with *SCN5A* variants transfected into the LP site, thereby allowing us to express the two SCN5A alleles at equivalent levels and assess dominant negative effects. These lines are referred to as LP-SB-SCN5A.

### Generation of Cell Lines

Cells were cultured at 37°C in humidified 95% air/5% CO_2_ incubator in “HEK media”: Dulbecco’s Eagle’s medium supplemented with 10% fetal bovine serum, 1% non-essential amino acids, and 1% penicillin/streptomycin. Stable integration of a WT *SCN5A* into LP-cells was achieved using an optimized SB transposon system (17) using the pSBbi-GN plasmid (a gift from Eric Kowarz, Addgene #60517), which contains SB transposon sequences for genomic integration flanking a promoter upstream of GFP and a second promoter upstream of a multiple cloning site (MCS) for expression of a gene of interest. A NotI restriction site was first cloned into the multiple cloning site using Gibson assembly (New England Biolabs). Then, WT *SCN5A* was cloned into the MCS by NotI digestion (New England Biolabs). Next, 1 ug of pSBbiGN-SCN5A and 100 ng of pCMV(CAT)T7-SB100, a plasmid expressing SB transposase (a gift from Zsuzsanna Izsvak, Addgene #34879), were cotransfected into the cells (18), using FuGENE 6 (Promega) following manufacturer’s instructions. At day 7 post-transfection, GFP+ cells were sorted by fluorescence-activated cell sorting (FACS), and individual colonies were picked and re-analyzed by analytical flow cytometry to identify clones expressing varying levels of GFP (and thus varying levels of Na_V_1.5). Clones were then tested by SyncroPatch automated patch clamping (see below) to identify a clone expressing an equal peak sodium current as results from typical integration of a single copy of wild-type Nav1.5 into the AttB/AttP landing pad.

For homozygous patch clamp experiments, LP cells were transfected with an AttB-SCN5A variant:IRES:mCherry-BlasticidinR plasmid and studied as previously described (14). For heterozygous patch clamp experiments, LP-SB-SCN5A cells were transfected using similar methods. For all cell lines, cells were transfected with FuGENE 6 or Lipofectamine 2000 following manufacturer’s suggested protocols using an AttB-containing SCN5A:IRES:mCherry:blasiticidinR plasmid and a plasmid bearing Bxb1 recombinase; cells underwent negative selection for 6 days with 1 ug/mL doxycycline (to induce promoter expression; Sigma), 100 ug/mL blasticidin S (to kill cells not expressing the blasticidin-resistant plasmid; Sigma), and 10 nM AP1903 (to kill un-integrated cells expressing the AP1903-sensitive caspase gene; MedChemExpress) in HEK media (15). At the end of selection, cells were assessed by analytical flow cytometry to assess percentage of mCherry-positive, BFP-negative cells (LP integration of *SCN5A* variant) and GFP-positive cells (SB integration of *SCN5A*).

### Automated Patch Clamping

Electrophysiology data were collected with the SyncroPatch 384PE automated patch clamping device (Nanion) using the same cell preparation and solutions as previously reported (14). Peak currents are reported at −20 mV after a 200 msec pulse from a resting potential of −120 mV; peak sodium current is presented as the mean of data obtained in ≥8 cells/variant (homozygous experiments) or ≥27 cells/variant (heterozygous experiments). Voltage of half activation, voltage of half inactivation, time of 50% recovery from inactivation, and late current at 200 ms were obtained using previously published protocols (14). As previously described, cells with values greater than 2.5 standard deviations from the mean were removed in an automated process (14). For these additional parameters, only variants with data collected from >10 cells were included.

### Case-control analysis

We performed case-control analyses to test the penetrance of different classes of variants. We used BrS case counts from a recent International BrS Genetics Consortium and putative controls from gnomAD; the frequency of these variants is presented in Supplementary Table 1 (12, 13). We use gnomAD as putative controls; although phenotypes are not available for gnomAD participants, the vast majority of these individuals should not have Brugada Syndrome. All gnomAD counts were taken from gnomAD v2.1.1 transcript ENST00000333535.4. A cut-off minor allele frequency of 2.5e-5 was used to designate ultra-rare variants, as previously suggested (19). To test the severity of each disease class (i.e., missense vs. indel vs. splice/frameshift/nonsense), we compared the relative number of cases versus controls by variant, drawing from the BrS consortium and gnomAD. Frameshift, splice, and nonsense variants at amino acid position > 1800 (post-transmembrane domain IV) were excluded due to the possibility that these variants may not be full LoF. We calculated the odds ratio associated with each variant class according to the formula (a/b)/(c/d), where a = BrS cases with variant, b = BrS cases without variant, c = gnomAD controls with variant, and d = gnomAD controls without variant. Since the allele number varied for different variants in gnomAD, the average allele number was calculated over all relevant mutation types (missense, frameshift, nonsense, and splice site) and divided by 2 to obtain a count of sequenced gnomAD participants to use in odds ratio calculations, following a previously published approach (12).

### Data Analysis

SyncroPatch 384PE data were analyzed as previously reported (14). Peak current densities were calculated by dividing peak current at −20 mV by cell capacitance. For homozygous experiments, peak current densities were normalized to peak current densities observed in cells expressing WT plasmid. For heterozygous experiments, peak current densities were normalized to that observed in LP-SB-SCN5A cells, *i.e*., those expressing a single WT allele. As described below, WT+WT cells displayed ~200% peak I_Na_ compared to LP-SB-SCN5A cells. Heterozygote (WT+variant) cells displaying <75% of peak I_Na_ compared to LP-SB-SCN5A cells were designated as exerting a dominant negative effect. Statistical comparisons were made using two-tailed Fisher’s exact tests, implemented in R Studio (version 1.3.1093).

### Structural Analysis

Na_V_1.5 variant locations were determined from UniProt (20). The structural model of human SCN5A (UniProtKB: Q14524-1, modeled residues: 30–440, 685–957, 1174–1887) was generated by homology modeling using the protein structure prediction software Rosetta (v.3.10) (21). The cryo-EM structure of human SCN9A bound with SCN1B and the Ig domain of SCN2B resolved to 3.2 Å (PDB: 6J8H) (22) were used as the primary templates while the cryo-EM structure of NavPaS from American Cockroach resolved to 2.6 Å (PDB: 6A95) (23) was used as a secondary template. The percent identity between the aligned positions of SCN9A and SCN5A sequences is 76.7%. While the percent identity between NavPaS and SCN5A was only moderate (45.6%), the N-terminal and C-terminal domains in the NavPaS structure were partially resolved, providing coordinates for modeling the corresponding domains of SCN5A. For further details, see our previous report (14). Recently, an experimental structure of SCN5A was determined using cryo-EM technique at a resolution of 3.3 Å (24). We note that the root-mean-square distance between our model and the experimental structure over all backbone atoms is 2.3 Å (Supplementary Figure 1), suggesting that our model is accurate while covering more residues than the experimental structure.

## Results

### Homozygous and Heterozygous Measurements of LoF Variants

We generated 37 LP-SCN5A stable lines (1 *SCN5A* allele expressed/line), each expressing LoF variants or the nonsense variant W822X (Figure 1, 2A and Table S1) (15, 16). Representative traces for WT and A735E are shown in Figure 2B. We recorded peak I_Na_ at −20 mV: 35/37 missense variants exhibited a peak current density <10% compared to WT (Figure 2C – only LoF shown; Supplementary Table 1). The remaining 2 variants (previously reported to be LoF) showed >10% peak current when compared to WT and were studied separately with other partial LoF variants. One LoF variant, R893C, was previously detected in patients with BrS but has not been previously assessed by patch clamping (2).

**Figure 2.**
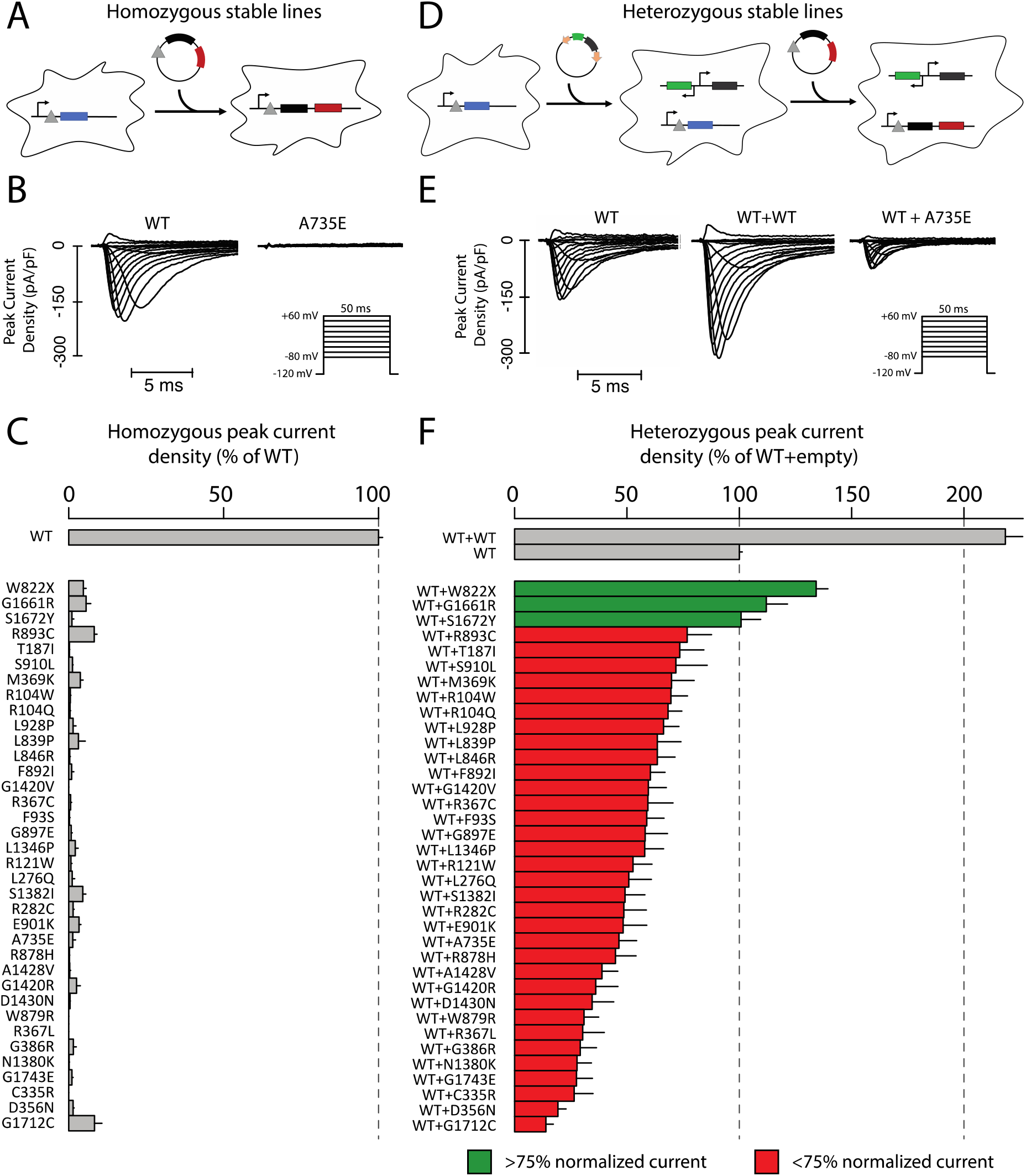
Measurement of loss-of-function homozygous and heterozygous peak current. **A**. Introduction of *SCN5A* variants into LP-*SCN5A* HEK cells. For full details see Figure 1. **B**. Representative raw peak current densities in a WT and A735E cell. Inset: voltage protocol used. **C.** Measurement of homozygous peak current density in 35 *SCN5A* missense variants and one nonsense variant (normalized to WT). Mean ± standard errors. 11-67 cells were studied per variant. **D.** Heterozygous LP-SB-*SCN5A* cell lines. For full details see Figure 1. **E.** Representative raw peak current densities in a single transfected WT, dually integrated WT+WT, and WT+A735E cell. **F.** Peak current density measurements for 35 *SCN5A* missense variants and one nonsense variant in expression with WT *SCN5A* (normalized to single WT). Mean ± standard errors. 27-164 cells were studied per variant.

We then tested each LoF variant in heterozygous expression (WT+variant) (Figure 1B and 2D). Figure 2E shows representative traces of cells expressing WT, WT+WT, and WT+A735E (an example dominant negative variant). Figure 2F presents peak I_Na_ for the same 35 LoF variants presented in Figure 2C. WT+WT cells expressed peak I_Na_ of 218.4±7.7% relative to WT alone in LP-SB-SCN5A cells, i.e., those expressing a single WT allele. By contrast, 32/35 of the WT+variant cell lines showed <75% peak I_Na_ compared to LP-SB-SCN5A cells, indicating a dominant negative effect. The heterozygous dominant negative variants displayed a gradient of effect, from 13.9±3.3% to 74.4±5.4% of WT alone. Two previously studied dominant negative variants, R104W and R121W (25), both also exhibited dominant negative effects in this study (69.6±7.3% and 52.7±8.4% of WT, respectively). While W822X, G1661R, S1672Y, and R893C had LoF peak currents in homozygous experiments, they did not exhibit a dominant negative effect (Table S1).

### Homozygous and Heterozygous Measurements of Partial LoF Variants

We also studied the prevalence of dominant negative effects in 15 partial LoF variants using LP-SB-SCN5A lines. We first confirmed that variant peak currents were 10%-50% compared to WT in homozygous expression with LP-SCN5A cells. (Figure 3A). The set of 15 variants included two variants (R282H and G1740R) previously reported to be LoF but measured as >10% peak I_Na_ in our system (26, 27). Figure 3B shows a gradient of I_Na_, with partial LoF variants showing a greater range of effect in heterozygous expression than those of LoF variants (24.7±5.6% to 231.6±10.8%). 6/15 partial LoF variants had a dominant negative effect whereas the remaining 9 variants all exceeded normalized WT peak current.

**Figure 3.**
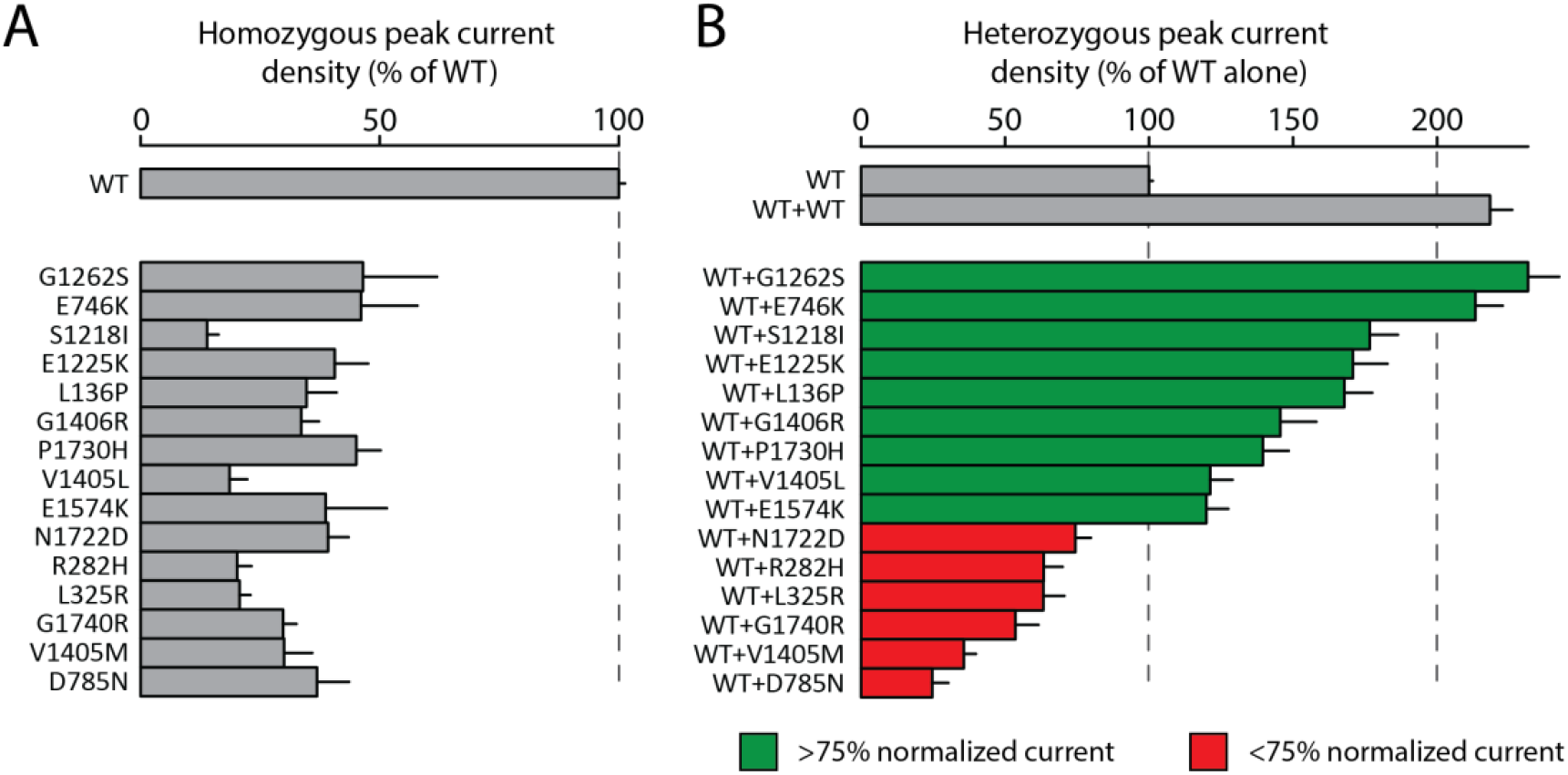
Some partial loss-of-function variants have a dominant negative effect. **A.** Measurement of homozygous peak current density in 15 partial LoF *SCN5A* variants (normalized to WT). Mean ± standard errors. 8-36 cells were studied per variant. **B.** Measurement of heterozygous peak current density in 15 partial LoF *SCN5A* variants (normalized to WT). Mean ± standard errors. 27-53 cells were studied per variant.

### Coupled gating in heterozygous expression

In addition to assessing peak sodium current, we also examined other parameters of channel function to measure the extent of coupled gating, a phenomenon where the LoF allele alters the gating properties of the WT allele. These parameters required additional experimental protocols and quality control filters, so these parameters were not comprehensively obtained in all variants studied; only variants with data from >10 qualifying cells are presented. We examined voltage of half activation among the missense variants investigated above (representative raw data shown in Figure 4A and 4B). 16/50 variants (14 LoF and 2 partial LoF) showed a >10 mV shift in the voltage of half activation, suggesting widespread coupled gating affecting this parameter (Figure 4C). We did not observe widespread changes for other parameters beyond voltage of half activation. No variants were shown to induce a shift in voltage of half inactivation >10 mV (Figure 4D). One variant (G1406R) had a 1.71-fold change in recovery from inactivation when compared to WT; the other 34 qualifying variants had <50% shifts in RFI (Figure 4E). No variants induced late current >1% when co-expressed with WT (Figure 4F). Due to the very low or absent peak currents in homozygous LoF variants, it was not feasible to assess parameters other than peak current in homozygous expression.

**Figure 4.**
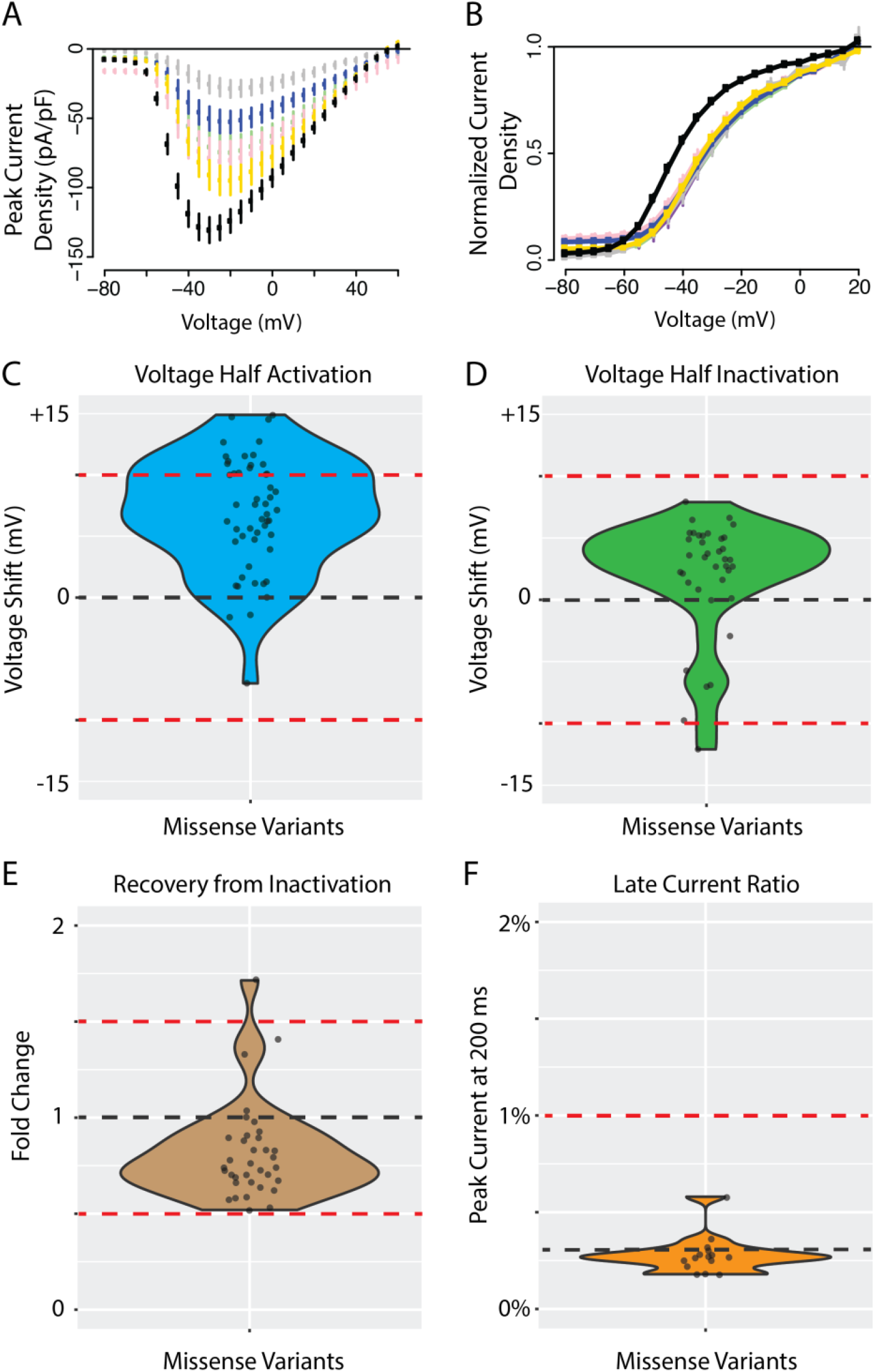
Additional channel parameters for missense variants in heterozygous expression. **A.** Current-voltage plot of WT (black) and 5 missense *SCN5A* variants with large shifts in voltage of half activation: A735E (light green), R121W (pink), D785N (grey), A1428V (blue), F892I (gold). **B.** Raw voltage half activation curve for WT and 5 missense *SCN5A* variants (variants and color same as in A). B-D) WT indicated with black line and abnormal cutoffs indicated with red lines. For B-D only variants with at least 10 qualifying cells meeting quality control criteria were analyzed. **C.** Voltage of half activation shift of all missense variants compared to WT. **D.** Voltage of half inactivation shift of all missense variants compared to WT. **E.** Time of 50% recovery from inactivation measured in fold change for all missense variants normalized to WT. **F.** Late current percentage (% of peak current) measured at 200 ms for all missense variants compared to WT.

### Elevated BrS Risk Among Dominant Negative Variants

Case and control counts of carriers of the dominant negative variants described above were interrogated using a published consortia of BrS cases (12) and gnomAD, a database of population variation that we considered to contain putative controls (13) (Figure 5A, Table S4). In Figure 5B we present the odds ratios (ratio of odds in BrS cohort:gnomAD). The LoF missense dominant negative variants had an odds ratio of 323 compared to 11.0 for missense, 24.2 for indel, and 118 for putative haploinsufficient variants (nonsense, splice, frameshift). Thus, the relative risk of dominant negative missense variants compared to haploinsufficient variants is 2.7 (Fisher’s exact test, p = 0.019). All categories were significantly enriched compared to all missense variants (Fisher’s exact test, p<0.05).

**Figure 5.**
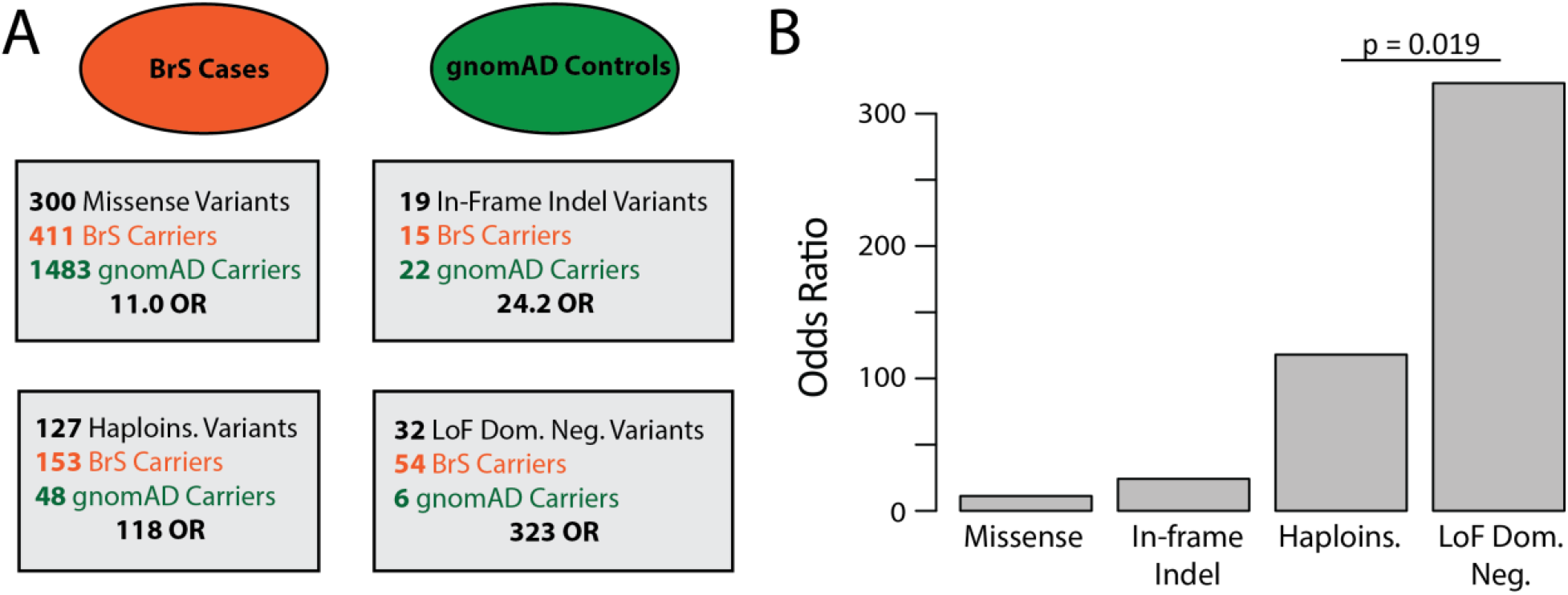
Case-Control analysis by variant class. **A.** Case-control breakdown by data source stratified by variant class. BrS cases are shown in red, with putative gnomAD controls shown in green. Haploins. Indicates nonsense, splice, and frameshift variants. Odds ratios are calculated for each variant class. **B.** Barplot of BrS odds ratios by variant class.

### Structural Distribution of Dominant Negative Variants

Dominant negative variants were present throughout the structured transmembrane regions of Na_V_1.5 and did not predominate in any single hotspot region (Figure 6A). Structural modeling further showed that dominant negative variants were distributed throughout the three-dimensional structure of Na_V_1.5, with apparent enrichment in the S5-S6 linker domains (Figure 6B and 6C).

**Figure 6.**
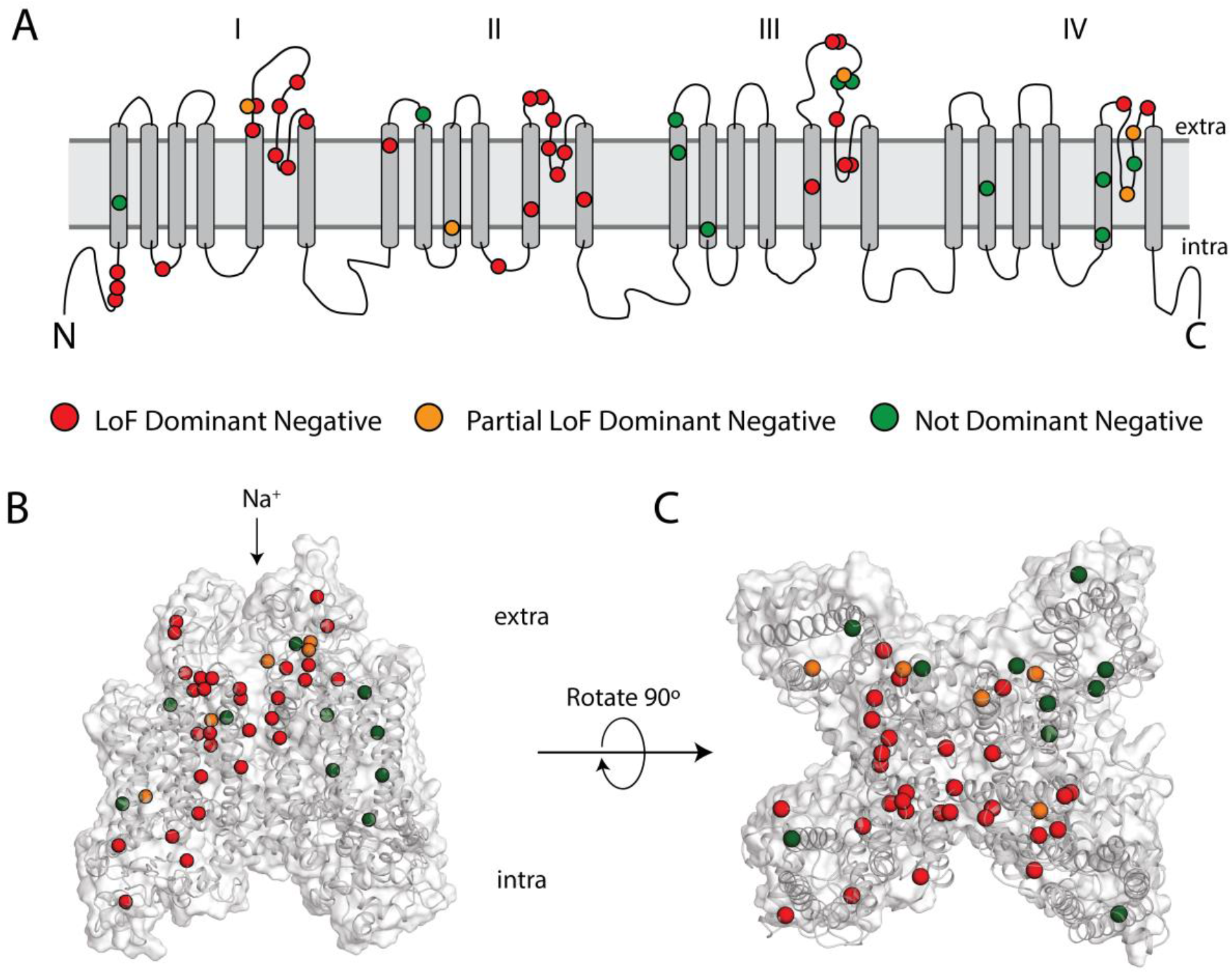
Structural distribution of dominant negative variants. **A.** Locations of dominant negative variants throughout Na_V_1.5 in 2D channel rendering. Red indicated LoF dominant negative, orange partial LoF dominant negative, and green non-dominant negative missense variants. Extra: extracellular, intra: intracellular. **B.** Side view of Na_V_1.5 protein with overlaid variant distribution. **C.** Top view of Na_V_1.5 protein with overlaid variant distribution.

## DISCUSSION

### Dominant Negative Effect Among Most Missense LoF *SCN5A* Variants

This study assessed the dominant negative properties of 50 LoF and partial LoF variants. A large majority of examined LoF variants (32/35) and some partial LoF variants (6/15) showed dominant negative behavior. Dominant negative effects are pervasive throughout biology, especially for multimeric proteins, and involve several distinct mechanisms to compromise WT function (28). In the case of *SCN5A*, the dominant negative effect has been posited to arise by both deficient trafficking to the membrane as well as coupled gating at the cell surface. One study showed that the variants R104W and R121W induced a dominant negative effect primarily through endoplasmic reticulum retention of WT protein due to interactions among the channel alpha-subunits (25). Follow up studies with extensive biochemical analyses showed that the dominant negative variant L325R acted through coupled gating at the cell surface (9).

Previous research suggested that the residues between 493 and 517 are critical for the dimerization and coupled gating of Na_V_1.5 at the cell surface, and another study found an enrichment of dominant negative variants at the N-terminus of the protein (8, 29). We did not observe an enrichment of dominant negative variants among these previously described residues, but rather a broader distribution of variants spanning the four transmembrane domains of the protein (Figure 6A-6C). Thus, dominant negative effects appear to be a general property of most LoF missense variants in *SCN5A*, independent of location within the protein. Particularly interesting are examples of disparate effects within close physical proximity, such as the partial LoF variants V1405M (35.7% peak current in heterozygous expression), V1405L (121% peak current), and G1406R (146% peak current).

In addition to decreased peak current, we observed that 16/50 variants also influenced voltage of half activation when measured in heterozygous expression with WT. This finding is consistent with the concept of coupled gating at the cell surface, and reflects the influence of the loss of function allele on properties of the WT allele of the protein, possibly through a multi-channel complex (9). These shifts in V1/2 activation in a loss of function direction combine with reduced peak currents to result in additional reduction of channel function in heterozygous expression. V_1/2_ activation was the only additional property that varied substantially from WT NaV1.5 activity, as we did not observe large differences in voltage of half inactivation, recovery from inactivation, or late current.

### Increased BrS Risk of Dominant Negative Variants

Previous work has established that homozygous peak current of *SCN5A* variants is the strongest *in vitro* electrophysiological predictor of each variant’s BrS risk (6, 30). Since dominant negative missense variants cause an especially low cellular peak current, we hypothesized that dominant negative variants would confer an especially high risk for BrS. Importantly, our expanded catalog of 38 dominant negative *SCN5A* variants enabled us for the first time to calculate cohort-based estimates of disease risk of this class of variants. Using gnomAD and a recently published cohort of BrS cases (12, 13), we demonstrated that dominant negative variants are highly overrepresented in cases vs controls when compared to other variant classes, with a striking odds ratio of 323 for dominant negative LoF missense variants. In contrast, other variant classes have lower odds ratios of 11 (all missense variants) or 118 (putative haploinsufficient frameshift/nonsense/splice site variants). Thus, the relative risk of BrS among dominant negative LoF missense variants compared to putative haploinsufficient variants is 2.7. Previous studies have shown that truncating and functionally inactive missense variants cause a more severe phenotype than partially active missense variants, but the penetrance of dominant negative variants had not yet been extensively studied (31). Our results indicate that the penetrance of dominant negative missense variants is higher than penetrance of other variant classes. One potential explanation for the different disease penetrance among variant classes is that nonsense mediated decay (NMD) removes aberrant transcripts for splice-altering and nonsense variants, preventing their interaction with WT Na_V_1.5. Given the data presented here, dominant negative missense variants should arouse high clinical suspicion for BrS risk when detected in patients.

### High-throughput Electrophysiological Assays to Study Dominant Negative Effects

High-throughput automated patch clamping has emerged as a tool for rapidly assessing functional consequences of ion channel genetic variation (32). This technique has been used to assess pathogenicity of variants in *KCNQ1* (33, 34), *SCN5A* (14), and *KCNH2* (35, 36). Here, we present the most extensive evaluation of heterozygous Na_V_1.5 expression to date using this platform, studying 51 variants with 27-164 cells per heterozygous measurement. Heterozygous measurements are already common for the cardiac potassium channels *KCNQ1* and *KCNH2*; this study suggests that heterozygous studies may also be necessary for LoF *SCN5A* variants in future studies. This work shows that high-throughput automated patch-clamp can help establish molecular mechanisms of disease.

### Mechanistic and Therapeutic Implications

The prevalence of widespread dominant negative effects among *SCN5A* variants not only gives insight into action potential pathophysiology, but also provides a lead for therapeutic development. The multifunctional regulatory protein 14-3-3 has been reported to be critical for mediating Na_V_1.5 dimerization, and an operative mechanism in select cases of the dominant negative effect (8). Indeed, difopein, an inhibitor of 14-3-3, (37) was shown to restore WT activity when co-expressed with dominant negative variants (8, 38). While targeting 14-3-3 may not be an appropriate therapeutic strategy given its role in myriad cellular processes, alternative mechanisms to promote selective degradation of aberrant channels and preserve WT function remain highly desirable. Emerging allele-specific siRNA or XNAzymes (39) strategies could ablate the dominant negative effect prior to the translation event. A gene therapy approach has recently been demonstrated for *KCNQ1*, and could be applied against dominant negative variants of BrS described here (40). Given the prevalence of the dominant negative phenomenon, and the high risk for BrS among carriers of these variants, there is a need for the development of novel therapeutic strategies by leveraging basic biological insights.

### Limitations

Results from heterologous expression in HEK293T cells may not fully recapitulate behavior in native cardiomyocytes in human hearts. In particular, contributions such as polygenic modifiers, as has been previously observed in BrS (41), may not be fully captured by this non-native system. Two common alternative splicing events impact SCN5A splicing (Q1077 deletion/insertion and fetal/adult exon 6); only the most common splice isoform in the adult heart was examined in this study. The gnomAD population database does not have available phenotypic information, so a small fraction of individuals included in gnomAD may in fact have BrS.

### Conclusions

Most LoF missense variants in SCN5A have a dominant negative effect. These missense dominant negative variants have a 2.7-fold increased risk of BrS when compared to putative haploinsufficient variants. These results may help refine prediction of BrS risk in dominant negative variant carriers.

## Acknowledgements

We thank Victoria Parikh for helpful discussions, Kenneth Matreyek for suppling the LP-negative HEK293 cell line, Eric Kowarz for supplying the pSBbi-GN plasmid, and Zsuzsanna Izsvak for supplying the pCMV(CAT)T7-SB100 plasmid. Flow Cytometry experiments were performed in the VMC Flow Cytometry Shared Resource. The VMC Flow Cytometry Shared Resource is supported by the Vanderbilt Ingram Cancer Center (P30 CA68485) and the Vanderbilt Digestive Disease Research Center(DK058404). SyncroPatch 384PE experiments were performed in the Vanderbilt High-Throughput Screening (HTS) Core Facility. The HTS Core receives support from the Vanderbilt Institute of Chemical Biology and the Vanderbilt Ingram Cancer Center (P30 CA68485).

---

**Table S1.** Variant currents and case-control counts.

| Variant | Homozygous |  |  | Heterozygous |  |  | gnomAD<br>Count | gnomAD<br>MAF | Walsh<br>Count |
| --- | --- | --- | --- | --- | --- | --- | --- | --- | --- |
|  | Peak<br>Current<br>Density | S.E. | Cells | Peak Current<br>Density | S.E. | Cells |  |  |  |
| WT | 100 | 1.3 | 2196 | 100 | 1.3 | 2196 | - | - | - |
| WT+WT | - | - | - | 218.4 | 7.7 | 199 | - | - | - |
| G1262S | 46.5 | 15.5 | 10 | 231.6 | 10.8 | 47 | 8 | 2.83E-05 | 3 |
| E746K | 46.1 | 11.8 | 9 | 213.3 | 9.5 | 45 | 6 | 2.14E-05 | 5 |
| S1218I | 13.9 | 2.4 | 19 | 176.6 | 9.8 | 47 | 1 | 4.02E-06 | 0 |
| E1225K | 40.6 | 7 | 19 | 170.8 | 12 | 43 | 1 | 4.01E-06 | 5 |
| L136P | 34.7 | 6.3 | 16 | 167.8 | 9.7 | 41 | 0 | 0 | 2 |
| G1406R | 33.6 | 3.7 | 18 | 145.6 | 12.5 | 35 | 0 | 0 | 3 |
| P1730H | 45.1 | 5.1 | 31 | 139.5 | 9 | 47 | 0 | 0 | 0 |
| W822X | 4.7 | 0.9 | 16 | 134.2 | 5.2 | 164 | 0 | 0 | 0 |
| V1405L | 18.6 | 3.7 | 15 | 121.2 | 7.8 | 53 | 0 | 0 | 4 |
| E1574K | 38.7 | 12.8 | 8 | 119.9 | 7.6 | 46 | 0 | 0 | 3 |
| G1661R | 5.6 | 1.5 | 19 | 112 | 9.4 | 44 | 0 | 0 | 3 |
| S1672Y | 1 | 0.6 | 18 | 100.8 | 8.7 | 47 | 0 | 0 | 1 |
| R893C | 8.2 | 0.9 | 48 | 76.8 | 10.8 | 52 | 3 | 1.06E-05 | 2 |
| N1722D | 39.2 | 4.3 | 26 | 74.4 | 5.4 | 43 | 0 | 0 | 1 |
| T187I | 0.2 | 0.1 | 42 | 73.5 | 10.7 | 39 | 0 | 0 | 1 |
| S910L | 1.2 | 0.2 | 19 | 71.8 | 13.9 | 35 | 1 | 3.99E-06 | 3 |
| M369K | 3.7 | 0.9 | 22 | 69.8 | 10.1 | 51 | 0 | 0 | 2 |
| R104W | 0.5 | 0.2 | 24 | 69.6 | 7.3 | 43 | 1 | 4.01E-06 | 3 |
| R104Q | 0.4 | 0.2 | 22 | 68.3 | 6.1 | 34 | 0 | 0 | 3 |
| L928P | 1.4 | 0.9 | 27 | 66.3 | 6.8 | 47 | 0 | 0 | 1 |
| L839P | 3.1 | 2.2 | 20 | 63.5 | 10.5 | 53 | 0 | 0 | 1 |
| L846R | 0.3 | 0.2 | 43 | 63.5 | 7.9 | 35 | 0 | 0 | 0 |
| R282H | 20.2 | 3 | 16 | 63.4 | 6.6 | 44 | 4 | 1.60E-05 | 8 |
| L325R | 20.7 | 2.3 | 36 | 63.3 | 7.3 | 49 | 0 | 0 | 0 |
| F892I | 0.9 | 0.7 | 23 | 60.4 | 6.5 | 51 | 0 | 0 | 1 |
| G1420V | 0 | 0 | 11 | 59.5 | 8 | 52 | 0 | 0 | 1 |
| R367C | 0.6 | 0.3 | 25 | 59.3 | 11.2 | 54 | 3 | 1.07E-05 | 3 |
| F93S | 0.2 | 0.2 | 15 | 58.8 | 7.7 | 53 | 0 | 0 | 1 |
| G897E | 0.8 | 0.3 | 16 | 58.1 | 9.9 | 38 | 0 | 0 | 0 |
| L1346P | 2.1 | 0.9 | 15 | 57.9 | 8.4 | 53 | 0 | 0 | 1 |
| G1740R | 29.8 | 2.8 | 20 | 53.6 | 8 | 27 | 0 | 0 | 1 |
| R121W | 0.7 | 0.3 | 40 | 52.7 | 8.4 | 36 | 0 | 0 | 3 |
| L276Q | 1.1 | 0.8 | 14 | 50.8 | 10.1 | 53 | 0 | 0 | 2 |
| S1382I | 4.5 | 1 | 29 | 49.1 | 8.9 | 47 | 0 | 0 | 1 |
| R282C | 1.4 | 0.3 | 67 | 48.6 | 10 | 55 | 0 | 0 | 2 |
| E901K | 3.3 | 0.6 | 16 | 48.3 | 10.5 | 46 | 0 | 0 | 6 |
| A735E | 1.3 | 0.9 | 12 | 46.4 | 7.9 | 39 | 0 | 0 | 0 |
| R878H | 0.2 | 0.1 | 38 | 44.9 | 9.1 | 39 | 0 | 0 | 3 |
| A1428V | 0.3 | 0.3 | 24 | 38.9 | 7 | 53 | 0 | 0 | 1 |
| G1420R | 2.5 | 1.2 | 16 | 36.1 | 9.9 | 50 | 0 | 0 | 2 |
| V1405M | 30 | 5.9 | 14 | 35.7 | 4.2 | 38 | 0 | 0 | 5 |
| D1430N | 0.4 | 0.1 | 57 | 34.5 | 9.6 | 28 | 0 | 0 | 0 |
| W879R | 0 | 0 | 43 | 30.9 | 6.5 | 46 | 0 | 0 | 0 |
| R367L | 0 | 0 | 39 | 30.3 | 9.6 | 46 | 0 | 0 | 1 |
| G386R | 1.5 | 0.9 | 11 | 29.2 | 7.2 | 52 | 0 | 0 | 0 |
| N1380K | 0.1 | 0.1 | 25 | 27.8 | 6.4 | 42 | 0 | 0 | 1 |
| G1743E | 1 | 0.4 | 11 | 27.5 | 7.1 | 37 | 0 | 0 | 5 |
| C335R | 0 | 0 | 24 | 26.5 | 8.4 | 27 | 0 | 0 | 1 |
| D785N | 36.9 | 6.7 | 27 | 24.7 | 5.6 | 33 | 0 | 0 | 0 |
| D356N | 1.4 | 0.3 | 16 | 19.3 | 3.6 | 45 | 1 | 4.02E-06 | 5 |
| G1712C | 8.3 | 2.4 | 17 | 13.9 | 3.3 | 38 | 0 | 0 | 0 |

**Table S2.**
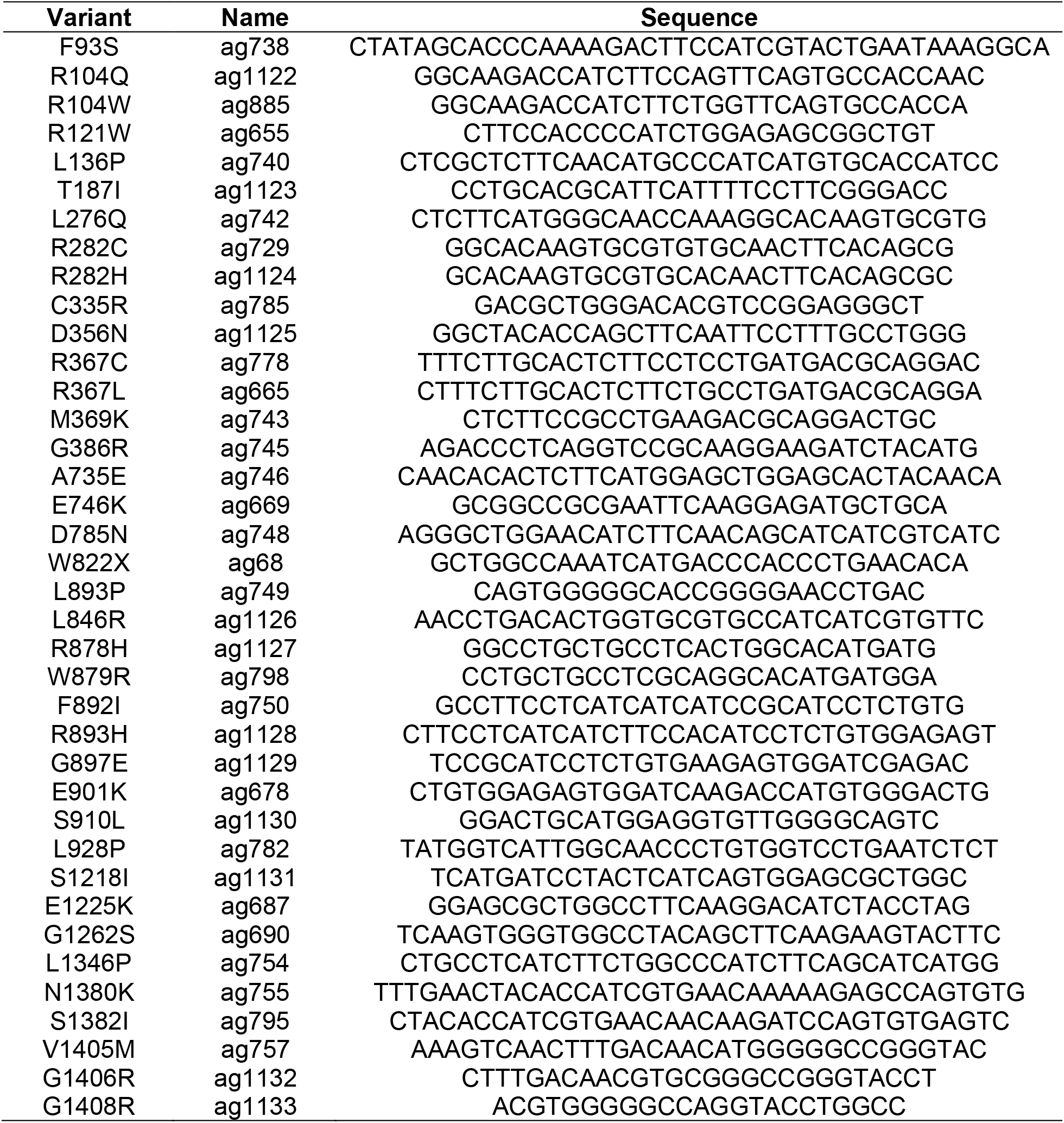
Primers used in this Study.

**Table S3.** Case-control analysis.

| **Class** | **# of**<br>**variants** | **BrS**<br>**cohort**<br>**count** | **gnomAD**<br>**count** | **gnomAD**<br>**AF** | **BrS :**<br>**gnomAD**<br>**ratio** | **Odds**<br>**Ratio** |
| --- | --- | --- | --- | --- | --- | --- |
| All missense | 300 | 411 | 1483 | 5.9e-3 | 0.28 | 11.0 |
| In-frame indel | 19 | 15 | 22 | 8.7e-5 | 0.68 | 24.2 |
| Frameshift+splice | 127 | 153 | 48 | 4.2e-4 | 3.19 | 118 |
| Missense LoF +<br>Dom. Neg. | 32 | 54 | 6 | 2.3e-5 | 9.0 | 323 |

**Figure S1.**
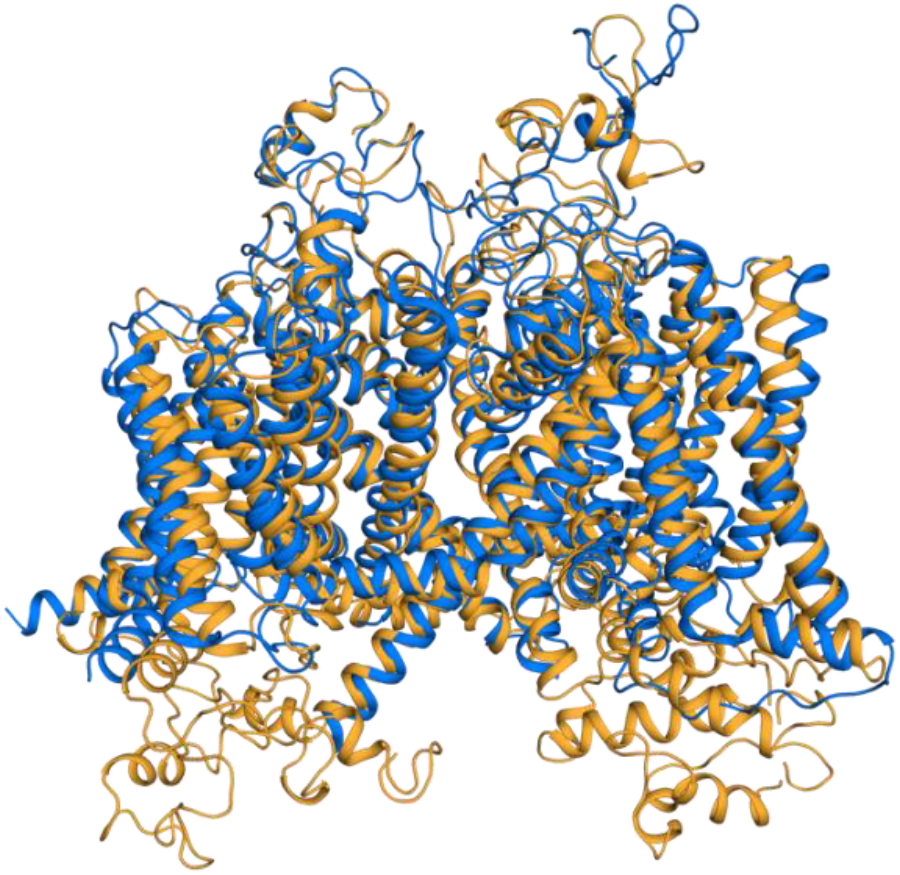
Overlay of our Nav1.5 structural model (light orange) with a recently determined cryo-EM structure of Nav1.5 (marine blue), demonstrating that our model is accurate while covering more intracellular residues than the experimental structure (24).

